# ANDROMEDA by Prosilico Software Successfully Predicts Human Clinical Pharmacokinetics of 300 Drugs Out of Reach for *In Vitro* Methods

**DOI:** 10.1101/2022.10.05.511015

**Authors:** Urban Fagerholm, Sven Hellberg, Jonathan Alvarsson, Ola Spjuth

**Affiliations:** Prosilico AB, Lännavägen 7, SE-141 45 Huddinge, Sweden; Department of Pharmaceutical Biosciences and Science for Life Laboratory, Uppsala University, Box 591, SE-751 24 Uppsala, Sweden

## Abstract

**Introduction:** *In vitro* measurements and predictions of human clinical pharmacokinetics (PK) are sometimes hindered and made impossible due to factors such as extensive binding to materials, low methodological sensitivity and large variability.

**Methods:** The objective was to find compounds out of reach for *in vitro* PK-methods and (if possible) predict corresponding human clinical estimates using the ANDROMEDA by Prosilico software. *In vitro* methods selected for the investigation were human microsomes and hepatocytes for measuring and predicting intrinsic hepatic metabolic clearance (CL_int_), Caco-2 and Ralph Russ canine kidney cells (RRCK) cells for measuring apparent intestinal permeability (P_app_) for prediction of fraction absorbed (f_a_), plasma for measurement and estimation of unbound fraction (f_u_), and water and buffers for measuring solubility (S) for prediction of *in vivo* dissolution potential (f_diss_).

**Results and Conclusion:** As many as 329 non-quantifiable *in vitro* PK-measurements for 300 compounds were found in the literature: 191 for CL_int_, 101 for P_app_, 11 for f_u_ and 26 for S. ANDROMEDA was successful in predicting all corresponding clinical PK-estimates for the selection of compounds with non-quantifiable *in vitro* PK, and predicted estimates (1.6-fold median prediction error; n=159) were generally in line with observed *in vivo* data and results/problems at *in vitro* laboratories. Thus, ANDROMEDA is applicable for predicting human clinical PK for compounds out of reach for laboratory methods.

## Introduction

Various *in vitro* methods are used to measure and screen pharmacokinetic (PK) properties of drug candidates, including human microsomes and hepatocytes for measuring and predicting intrinsic hepatic metabolic clearance (CL_int_), Caco-2, Madin Darby canine kidney (MDCK) and Ralph Russ canine kidney (RRCK) cells for measuring apparent intestinal permeability (P_app_) for prediction of fraction absorbed (f_a_), plasma for measurement and estimation of unbound fraction (f_u_), and water and buffers for measuring solubility (S) for prediction of *in vivo* dissolution potential (f_diss_).

*In vitro* measurements and *in vivo* predictions are commonly hindered and made impossible due to extensive binding to materials, low methodological sensitivity and large variability between laboratories and method set-ups. For example, the *in vitro* PK for about every other compound could not be quantified with human microsomes and hepatocytes (CL_int_<limit of quantification (LOQ)) and Caco-2 cells (low recovery) in studies by Stringer et al. (2008) and Skolnik et al. (2010), respectively. This has recently also been investigated and shown by Fagerholm et al. (2022a-c).

Lack of preclinical PK data/information could jeopardize drug discovery and development, for example, causing systemic underexposures (and many dosing arms) or unwanted overexposures (side-effects) in first clinical trials with candidate drugs, resulting in selection of compounds with inadequate *in vivo* PK and making optimization of PK-properties difficult/impossible. Thus, it is important to improve and develop new methodologies so that useful PK-data can be produced for compounds that are out of reach for the conventional *in vitro* methods.

The ANDROMEDA software by Prosilico for prediction, simulation and optimization of human clinical PK has been applied and validated in many studies (Fagerholm et al. 2021a,b and 2022a,d,e). It has been shown to outperform (higher accuracy) laboratory methods in 13 comparisons out of 17 (76 %; on par in 12 % of cases) and to have a wider prediction domain (with ability to predict human clinical PK for compounds with good metabolical stability (low CL_int_), extrahepatic elimination and low P_app_, S and recovery) (Fagerholm et al. 2022a,e)).

The objective of this investigation was to - via literature search - find compounds out of reach for *in vitro* PK-methods (human microsome and hepatocyte CL_int_, P_app_, plasma f_u_ and aqueous S) and then to (if possible) predict corresponding human clinical estimates using ANDROMEDA.

## Materials & Methods

The literature was searched for compounds with non-quantifiable human microsome and hepatocyte CL_int_ and P_app_, uncertain f_u_ (below a certain limit) and reported insolubility in water (including lack of S-data). Useful data were found in the following references: microsome and hepatocvte CL_int_ (Stringer et al. 2008; Sohlenius-Sternbeck et al. 2010; Obach 1999; Riley et al. 2005; McGinnity et al. 2007; Keefer et al. 2023; own non-published data), Caco-2 and RRCK P_app_ (Irvine et al. 1999; Fagerholm and Björnsson 2005; Sköld et al. 2006; Alsenz and Haenel 2003; Keefer et al. 2023), f_u_ (Fagerholm el al. 2021c; FDA drug labels) and S (Pham-The et al. 2013; Jiménez et al. 2022; Newby et al. 2023; Prosilico data bank).

ANDROMEDA software by Prosilico for prediction, simulation and optimization of human clinical PK) was applied to predict the main parameters for the study: *in vivo* CL_int_, f_a_ (permeability-based f_a_), f_u_, and f_diss_ (at a default oral dose of 50 mg).

## Results, Discussion & Conclusion

In total, 329 non-quantifiable *in vitro* PK-measurements for 300 compounds were found: 191 for CL_int_, 101 for P_app_, 11 for f_u_ and 26 for S (Table 1).

**Table 1.**
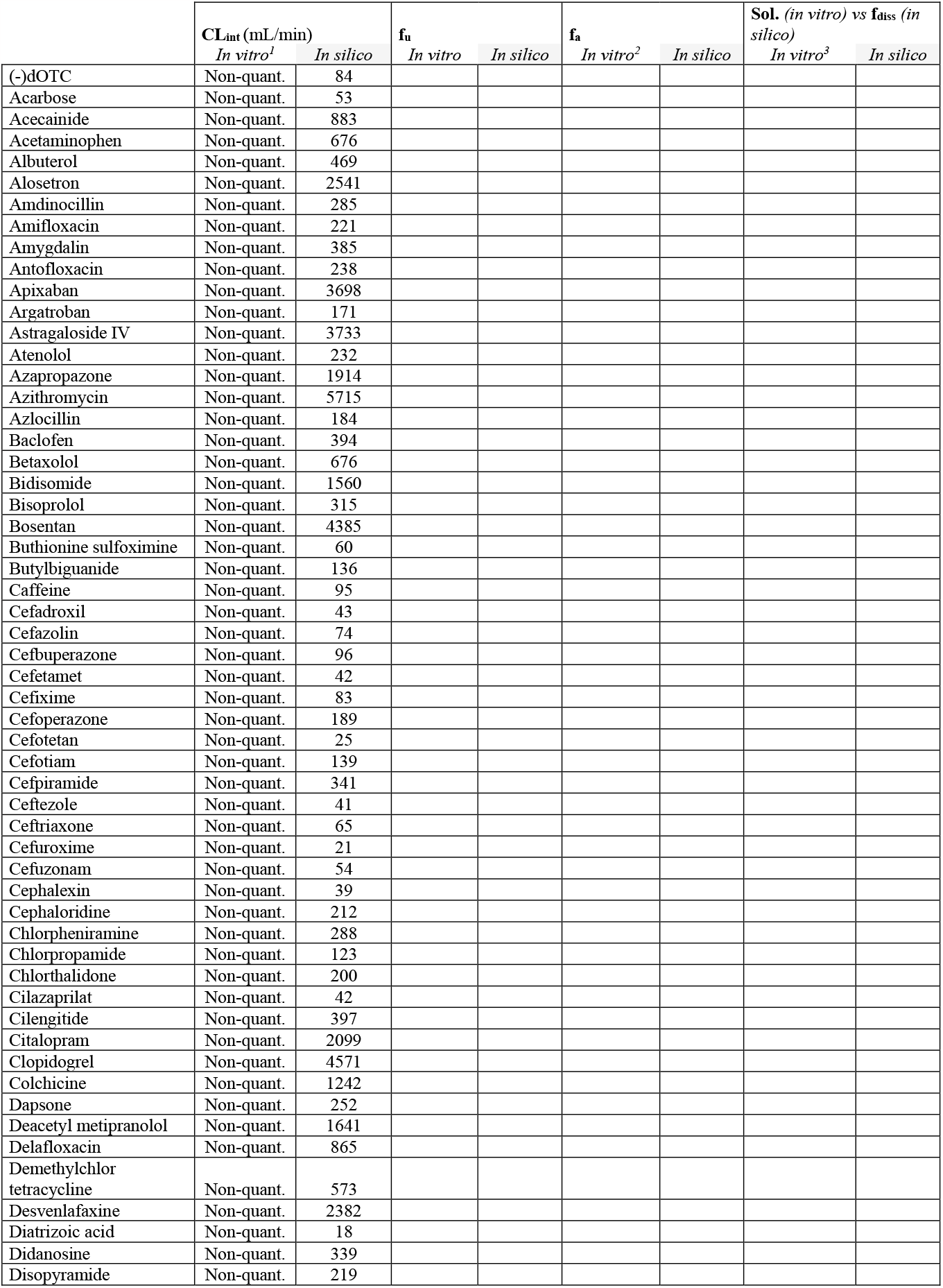
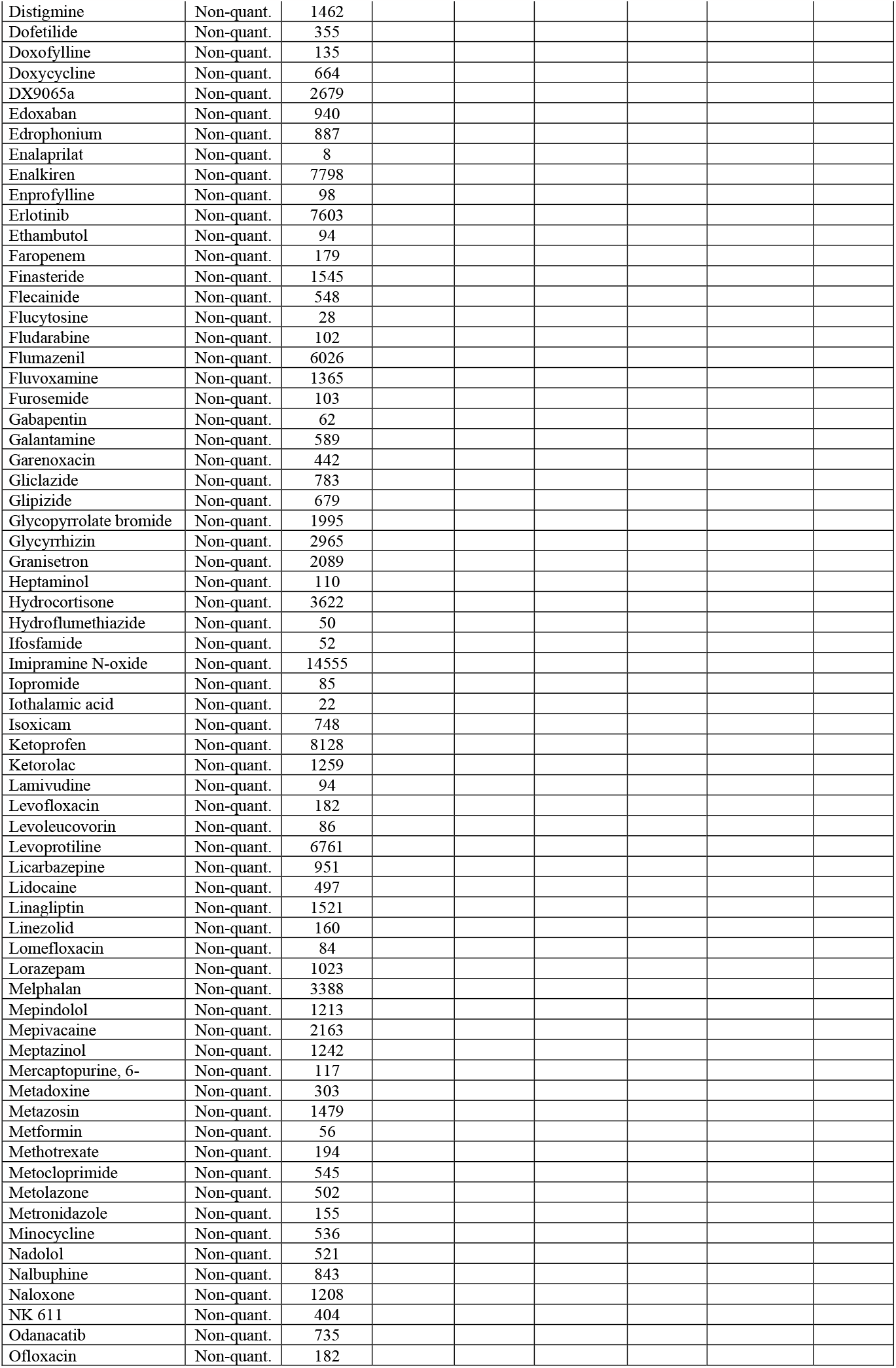
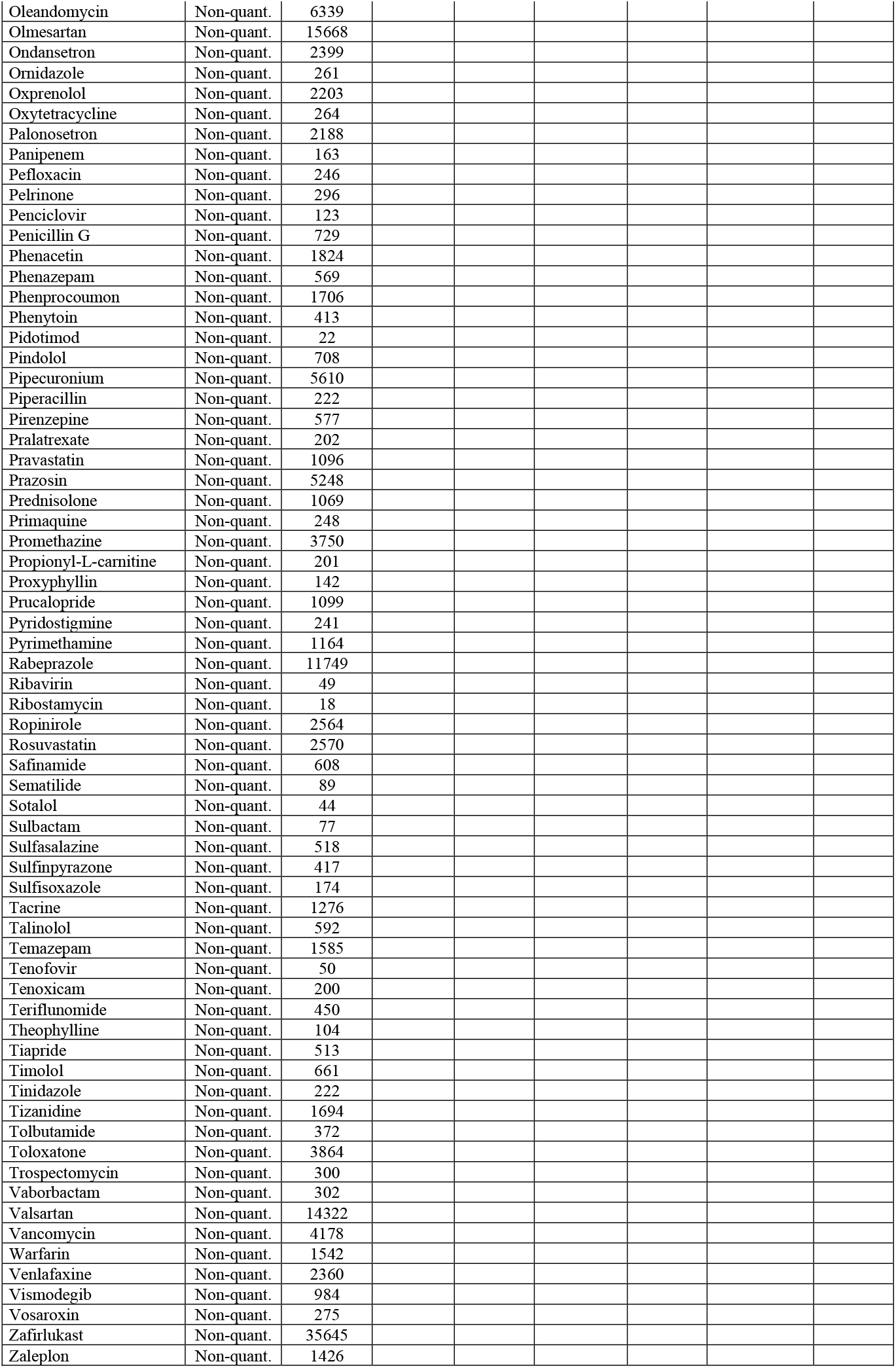
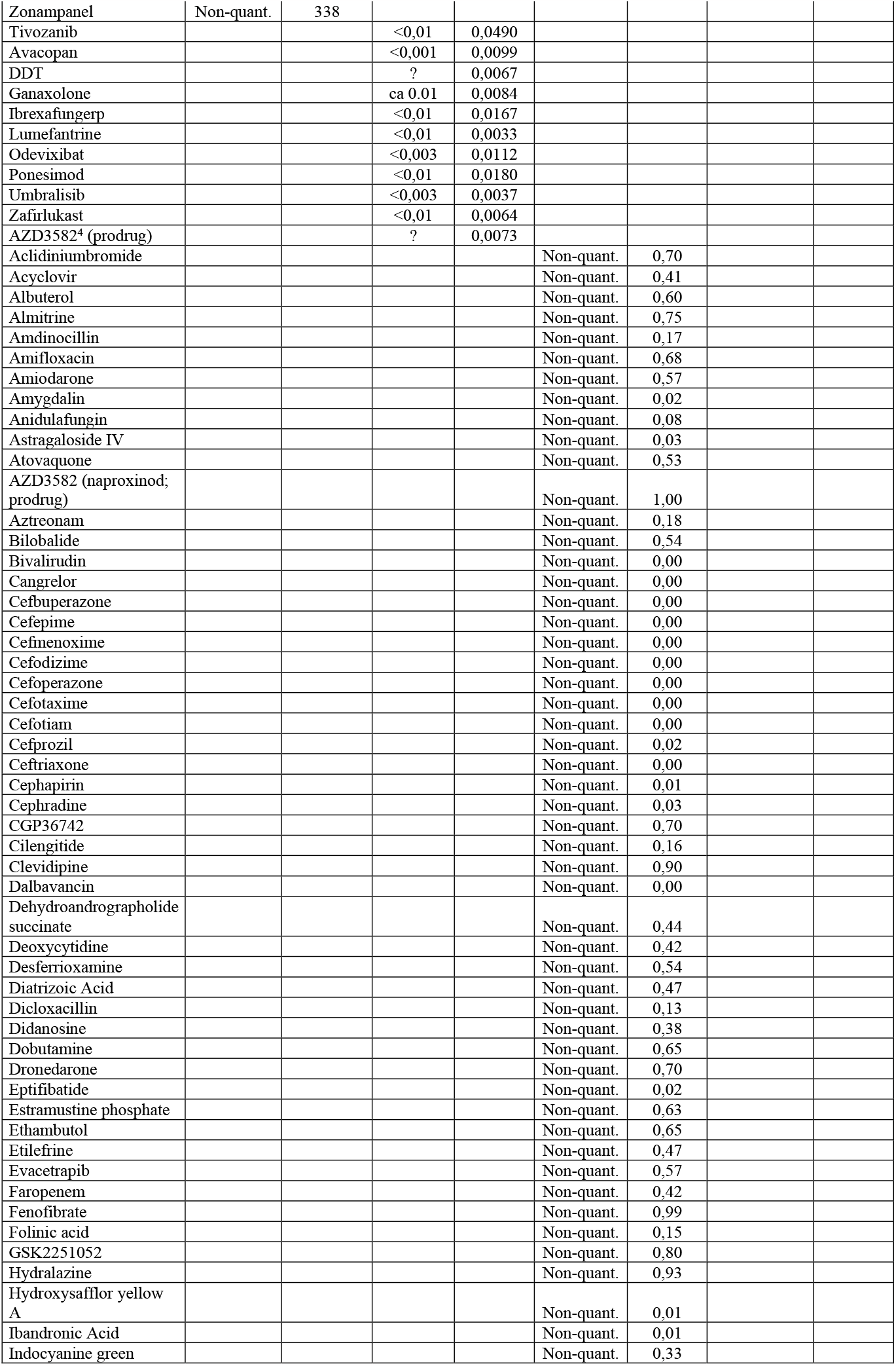
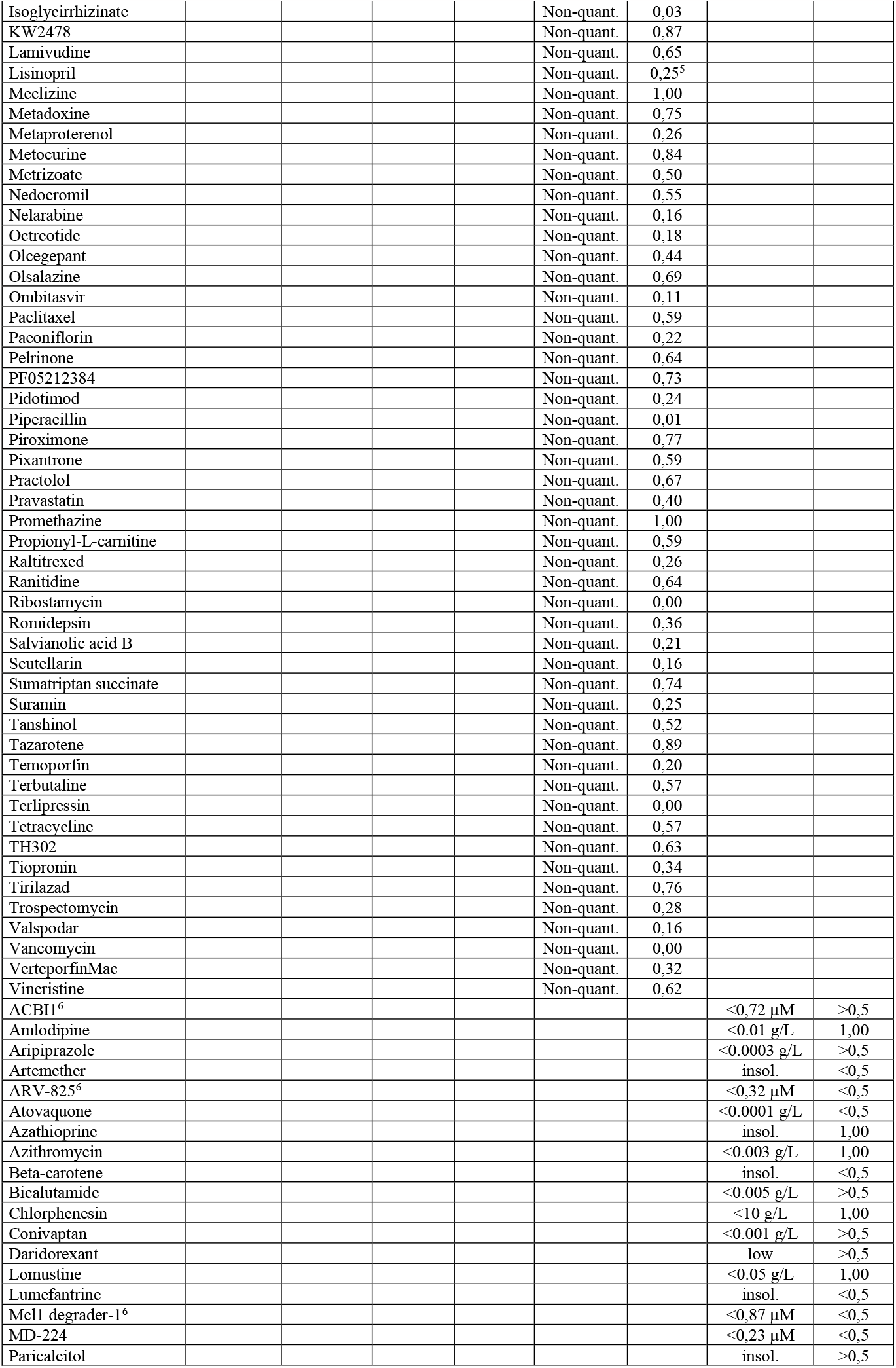
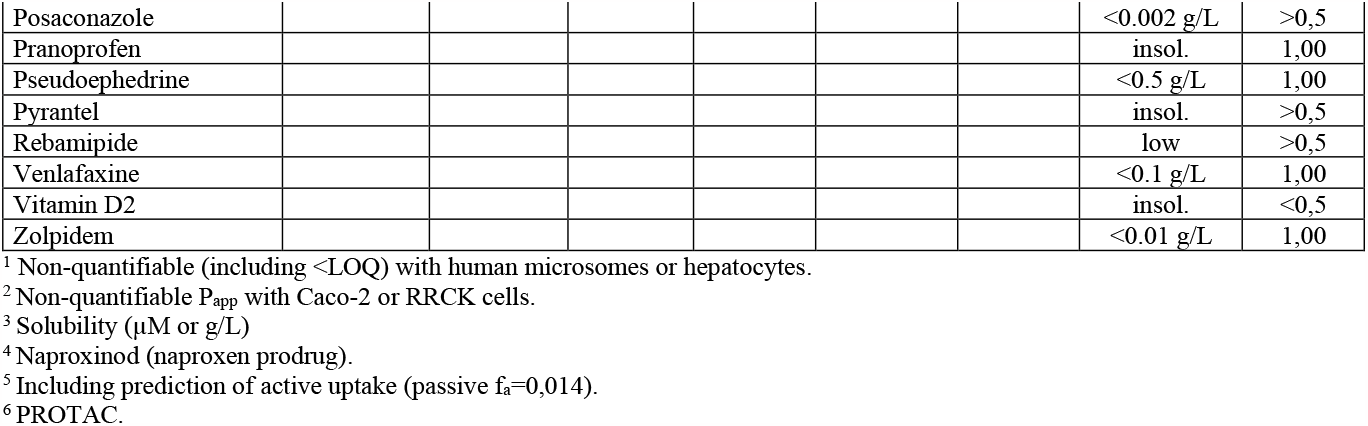
300 compounds with non-quantifiable *in vitro* PK and corresponding *in silico* predicted estimates.

The minimum, median and maximum predicted CL_int_ were 8, 442 and 35645 mL/min, respectively. 78 (60 %) of the compounds with *in vitro* CL_int_<LOQ had an *in vivo* CL_int_<500 mL/min, a limit that is slightly below the median *in vivo* CL_int_ for drugs (738 mL/min) and about 100-fold higher that lowest reported clinical values (Fagerholm 2022b,c; Varma et al. 2010). 55% of CL_int_-predictions had an error of <2-fold (n=130), which in consistent with previous prediction results with ANDROMEDA (Fagerholm et al. 2022a,e). For these low CL_int_-compounds it is common with contribution by elimination via excretion (Fagerholm 2022b). Thus, human microsomes and hepatocytes are generally insufficient for measuring and predicting elimination of such compounds. Validated prediction models for renal and biliary excretion are included in ANDROMEDA.

The predicted and observed f_a_ for the compounds lacking P_app_-data due to low recovery or sensitivity limitations were ca 0 to 1.00. The median-fold prediction error for f_a_ was 1.4-fold (n=29). For the hydrolysis sensitive naproxen-prodrug AZD3582 (naproxinod) an intrinsic f_a_ (in case of absence of gastrointestinal degradation) of 1.00 was predicted. The passive f_a_ of actively absorbed ACE-lisinopril was predicted to 0.01. Considering the active component a f_a_ of 0.25 was predicted.

The overall median prediction error for CL_int_ and f_a_ was 1.57-fold.

Predicted f_u_-values ranged between 0.0033 and 0.049 and were generally close to reported maximum f_u_-estimates (<0.001 to <0.01). In 3 cases predicted estimates were lower than measured upper limits (<X), and in 6 cases they were above them.

For compounds with unmeasurable S, f_diss_ at the 50 mg oral dose level ranged between 0.01 (beta-carotene) and 1.00, with 8 (31 %) of compounds with f_diss_<0.5. Incomplete f_diss_ was predicted for 17 (65 %) of the 26 compounds. As described in recent papers, there is very weak correlation between the clinical solubility/dissolution parameter f_diss_ and *in vitro* S (Fagerholm 2022b,c).

In conclusion, ANDROMEDA was successful in predicting all the corresponding clinical PK-estimates for the selection of 300 compounds with non-quantifiable *in vitro* PK, and predicted estimates were generally in line with observed *in vivo* data and results/problems at *in vitro* laboratories. Thus, this software is applicable for predicting human clinical PK for compounds out of reach for laboratory methods.

